# Efficient Fluorescence-Based Localization Technique for Real-Time Tracking Endophytes Route in Host-Plants Colonization

**DOI:** 10.1101/2021.09.30.462628

**Authors:** Christine A. Ondzighi-Assoume, Bandana Bhusal, Adam Traore, Wilson K. Ouma, Margaret Mmbaga, Ethan Swiggart

## Abstract

Bacterial isolates that enhance plant growth and suppress plant pathogens growth are essential tools for reducing pesticide applications in plant production systems. The objectives of this study were to develop a reliable fluorescence-based technique for labeling bacterial isolates selected as biological control agents (BCAs) to allow their direct tracking in the host-plant interactions, understand the BCA localization within their host plants, and the route of plant colonization. Objectives were achieved by developing competent BCAs transformed with two plasmids, pBSU101 and pANIC-10A, containing reporter genes *eGFP* and *pporRFP*, respectively. Our results revealed that the plasmid-mediated transformation efficiencies of antibiotic-resistant competent BCAs identified as PSL, IMC8, and PS were up 84%. Fluorescent BCA-tagged reporter genes were associated with roots and hypocotyls but not with leaves or stems and were confirmed by fluoresence microscopy and PCR analyses in colonized Arabidopsis and sorghum. This fluorescence-based technique’s high resolution and reproducibility make it a platform-independent system that allows tracking of BCAs spatially within plant tissues, enabling assessment of the movement and niches of BCAs within colonized plants. Steps for producing and transforming competent fluorescent BCAs, as well as the inoculation of plants with transformed BCAs, localization, and confirmation of fluorescent BCAs through fluorescence imaging and PCR, are provided in this manuscript. This study features host-plant interactions and subsequently biological and physiological mechanisms implicated in these interactions. The maximum time to complete all the steps of this protocol is approximately three months.

**SENTENCE SUMMARY:** We describe a novel fluorescence localization technique as a powerful tool to directly visualize and determine the route *in-situ* of BCAs in host-plants interaction. The study features the host-plant interactions, biological and physiological responses implicated.

## INTRODUCTION

Principal types of microbe-plant interactions such as parasitism, competition, commensalism, and mutualism are constantly integrated parts of our ecosystem. Commensalism and mutualism are the more common interactions, where either one or both species benefit from the relationship, respectively (Campbell, 1995). Plants interact with various microorganisms in their natural surroundings, with some organisms colonizing their internal tissues as endophytes without harming their host plants (Petrini, 1988). Among these endophytes, bacteria in the genus *Bacillus* are known to be the most important bacterial antagonists comprising biological control agents (BCAs) whose numbers and uses are rapidly increasing (Shafi et al. 2017, Verschuere et al. 2000; Irabor and Mmbaga 2017; Bhusal and Mmbaga, 2020; Maheshwari et al. 2021). Endophytes have gained popularity as practical tools for sustainable and environmentally friendly approaches in modern agriculture, serving as biopesticides, biofertilizers, or phytostimulants (Bonaldi et al. 2015). The ability of endophytes to inhibit or antagonize plant pathogens is through different mechanisms, including the production of volatile or diffusible antimicrobial compounds, enzymes for degrading phytopathogen cell walls, antibiotics, and plant growth-promoting compounds, such as growth hormones, ammonia, phosphate solubilizing as well as siderophores for nutrients acquisition (Loaces et al. 2011; D’Alessandro et al. 2014; Khan et al. 2016; Li et al. 2016; Maheshwari et al. 2021).

Several endophyte BCAs have been isolated and identified in our earlier studies (Mmbaga et al. 2008; Joshua and Mmbaga 2019). In these studies, BCAs isolated from flowering dogwood, papaya, pepper and snap beans were shown to exhibit biocontrol activities against fungal diseases (Mmbaga et al. 2008, Rotich, 2015; Mmbaga et al. 2016; Irabor and Mmbaga, 2017; Joshua, 2017; Mmbaga et al. 2018a, 2018b) and actively produce secondary metabolites for antagonism of plant pathogens (Raaijmakers et al. 2002; Compant et al. 2005). They were also shown to produce various antimicrobial secondary metabolites, which can inhibit the growth of fungal pathogens. However, the biological process and mechanism by which Bacillus species control plant pathogens is still unclear because of the lack of genetic manipulation system (Yang et al. 2014). Thus, for these advantages, it is crucial to unravel the biotechnological potential of these BCAs, especially for understanding their proper applications (Villarino et al. 2018).

Biological control agents are emerging as essential tools in agriculture, especially in plant-microbe interactions, mainly due to their ability to control pathogen infection, induce resistance, and increase growth rate. BCAs potential as a useful agricultural tool depends heavily on multiple environmental factors that vary their performance in laboratory, greenhouse, and field conditions. Previous works showed that antibiotic selection is a critical marker for studying root colonization by introduced beneficial microorganisms (Gamalero et al. 2003). However, molecular techniques in conjunction with microscopy are among the most promising techniques for identifying microorganisms inside plant hosts. Colonization and localization of endophytic microorganisms can be determined with the help of several fluorescent markers (Lu et al. 2004; Cao et al. 2011; Krzyzanowska et al. 2012). Fluorescent proteins are an essential tool for identifying prokaryotic organisms (Parker and Bermudez, 1997; Valdivia and Falkow, 1997; Bumann 2001; Poschet et al. 2001). The use of fluorescent proteins (GFP and RFP) tagged to bacteria is easier to visualize under a microscope within a cell.

Additionally, its use for colocalization studies is less time-consuming in comparison to the use of dyes or other reporters (Bloemberg, 2007). Although several previous studies (Lu et al. 2004; Bolwerk et al. 2005; Grunewaldt-Stöcker et al. 2007) have demonstrated that fluorescence labeling of endophytes with fluorescence tags contributed to study the mechanisms of host-microbe interaction, the localization technique of these bacteria remains unfeasible. Transformed bacterial populations’ stability is influenced by plasmid-mediated changing environmental conditions (Smalla et al. 2015).

Despite using these promising techniques, the colonization and exact niches/locations of the BCAs inside the host-plant remain to be determined. In this context, several questions remain to be addressed. How and where do BCAs physically interact, traffic, and localize within their plant hosts? In order to address these questions, our study focused on developing a sustainable and reliable fluorescence-based localization technology for BCAs to track their itinerary in real-time within the host-plant organs directly. Thus, four objectives were investigated, including (i) developing competent BCA strains capable of genetic transformation, (ii) developing reliable plasmid-mediated transformation methods for BCAs, (iii) developing and implementing inoculation and growth procedures to study host-plants interaction, and (iv) determining BCA spatiotemporal niches/locations of BCAs in transformed plants using molecular genetics and microscopy techniques.

As we demonstrate here, this method can be applied to observe single-cell or bulk bacteria behavior within plant cells and organs during their colonization in host-plant interactions. It is also applicable to study bacteria endophytes and pathogens or pest interactions in host-plant infection, which have until recently primarily relied on static imaging or ex-vivo models for important spatial and temporal information. In this study, we use an upright fluorescence microscope system with a tunable fluorescence filter set. Ideally, scanning microscopes with laser multiphoton will be more suitable to live-imaging and capture high-resolution 3-D images of single cells type in dense tissues. A high-resolution fluorescence stereomicroscope will be suitable as well for big specimens. The use of dual localization (GFP-RFP) implemented on a dual-laser multiphoton microscope is a relatively straightforward strategy that will benefit others planning to co-image BCA-BCA, BCA-pathogens, or BCA-pathogen-plant interactions in dense tissues.

Biotechnology, molecular biology, microscopy, and image analysis skills are required to perform this procedure and analyze the resulting data. Skills in cell and tissue culture are also beneficial. It takes approximately three months to become proficient in this technique with training. Specialized imaging equipment is also needed; the most crucial equipment is an upright epifluorescence microscope, an upright confocal microscope, or a fluorescence stereomicroscope.

The flow chart and timeline of the procedure comprise five main steps: 1). Bacteria BCA strains source, growth, and maintenance conditions: Four selected Bacillus species utilized as biological materials listed and featured in Table 1 were used for this study. The process of growth and maintenance of bacterial strains took 2-3 days. 2). Generation of competent BCA cells by chemical induction: The competent BCA cells were generated with two high-level methods: (i) The TRIS-HCl-induced competency method, 10 mM Tris-HCl, pH 7.5 as described by Hofgen and Willmitzer (1988), and (ii) The Calcium Chloride-induced method, 0.1 M Calcium Chloride method adapted from Chang et al. (2017). The production of competent BCA cells was done for 2-3 days. 3). Plasmid-mediated heat-shot transformation of BCA cell suspensions. The transformation procedure was conducted using a method developed in our laboratory modified from previous protocols (Aymanns et al. 2011; Ondzighi-Assoume et al. 2019). The transformation and generation of transformation efficiencies data of competent BCA cells were done in 2-3 days. 4). Host-plant interaction and growth conditions: BCA-mediated colonization in plants. For the colonization and monitoring growth conditions of fluorescent BCAs in plants, two different plants *Arabidopsis thaliana* genotype Columbia (Col-0) and sweet sorghum variety Topper, 76-6, were used in this study. This procedure took 9-16 days. All steps in this protocol were performed aseptically in the biosafety cabinet logic A2 surface decontaminated with 70% ethanol. 5). The analysis includes (i) confirmation of the transformation of competent BCAs through both plasmid and genomic DNA isolations and PCR assay, (ii) fluorescence microscopy analysis with the visualization of fluorescent BCAs population and their spatial-temporal localization in colonized plants, and (iii) statistical analysis. All sub-steps in this analysis were completed in about two months. The maximum time to complete all the steps of this protocol is about three months.

**Table 1.** Transformation efficiencies of the pBSU101-mediated transformation of BCA cells. The numbers represent the efficiencies of the plasmid pBSU101 transformation of BCA strain IMC8, PRT, PS, and PSL cultures. The CaCl_2_- or TRIS-induced competent BCA cells were transformed simultaneously with the final concentration of inoculum OD_600_ = 0.5. The transformation efficiency was evaluated by scoring growing spectinomycin (*SPEC*^*R*^)-resistant colony-forming unit (CFU). The data depicted in the table corresponds to the mean ± SD of two replications of transformation events (n = 3 plates scored per transformation event for each BCA strain).

|  |  | pBSU101 |  |  |  |  |  |
| --- | --- | --- | --- | --- | --- | --- | --- |
| BCA strain | Selection antibiotic | CaCl<sub>2</sub> |  |  | TRIS |  |  |
|  |  | Total no. of CFU grown | Total no. of Resistant CFU | Percentage efficiency (%) | Total no. of CFU grown | Total no. of Resistant CFU | Percentage efficiency (%) |
| IMC8 | *SPEC<sup>R</sup>* | 659.67±51.03 | 382.33±19.66 | 58.36±7.78<sup>b</sup> | 765.33±33.26 | 393±17.06 | 51.48±4.38<sup>b</sup> |
| PRT | *SPEC<sup>R</sup>* | 742.33±51.25 | 408.67±84.5 | 54.86±8.97<sup>b</sup> | 193±27.84 | 108±17.35 | 57.57±16.74<sup>ab</sup> |
| PS | *SPEC<sup>R</sup>* | 120.33±22.19 | 82±11.14 | 69.09±11.39<sup>ab</sup> | 52±22.34 | 35.67±7.37 | 73.63±20.85<sup>ab</sup> |
| PSL | *SPEC<sup>R</sup>* | 102±10.15 | 4.67±3.79 | 5.43±2.39<sup>c</sup> | 146.67±14.05 | 123.33±18.58 | 83.89±7.41<sup>a</sup> |

## RESULTS

### Performance of BCAs in the Competency: Growth Characteristics of Produced Competent BCA Strains

For the performance of BCA strains in the competency, growth characteristics of competent BCAs produced were assayed using a spectrophotometer. To assess the influence of either TRIS-or CaCl2-induced competency on BCA strains IMC8, PRT, PS, and PSL, frozen bacteria with 0.5 OD_600_ at the mid-log phase of growth were grown on LB solid medium first and then sub-cultured in LB liquid at different time points (1hr, 4 hrs, and overnight). We found that the strain of bacteria and competency method used was important to achieve reproducible competency of BCA strains. TRIS-treated and CaCl2-treated BCAs IMC8, PRT, PS, and PSL strains grew well, displaying titers varied from 0.1±0.18 to 0.27±0.39 OD_600_ by 1 hr of culture, up to 0.2±0.22 to 0.67±0.59 OD_600_ by 4 hrs and reaching up to 0.5±0.37 to 1.03±1.11 OD_600_ by overnight (Fig. 1a, b, c). Either TRIS-treated or CaCl2-treated BCAs cell cultures displayed similar densities of cells over each time of culture compared to H_2_O-treated BCAs (CTR); thus, indicating that the competency did not impact the growth of BCA strains. Additionally, when optical densities were compared, we determined that BCAs growth for 1 hr would be appropriate inoculums for further transformation assays. Similarly, the influence of cold/freezing on the growth of stored competent BCAs were evaluated. We found that ODs of either frozen CaCl2-treated or TRIS-treated BCAs were significantly elevated for CaCl2-treated and TRIS-treaded BCA IMC8 compared to non-treated BCA IMC8 for an overnight culture, with up to 1.3- and 1.7-fold increase, respectively. However, no significant differences in optical densities were observed for either CaCl2-treated or TRIS-treated BCA PRT, PS, and PSL compared to controls (CTR) for an overnight culture (Supplementary Fig. 1a). These observations suggest that the cold or freezing did not negatively impact the growth of competent BCA cells.

**Figure 1.**
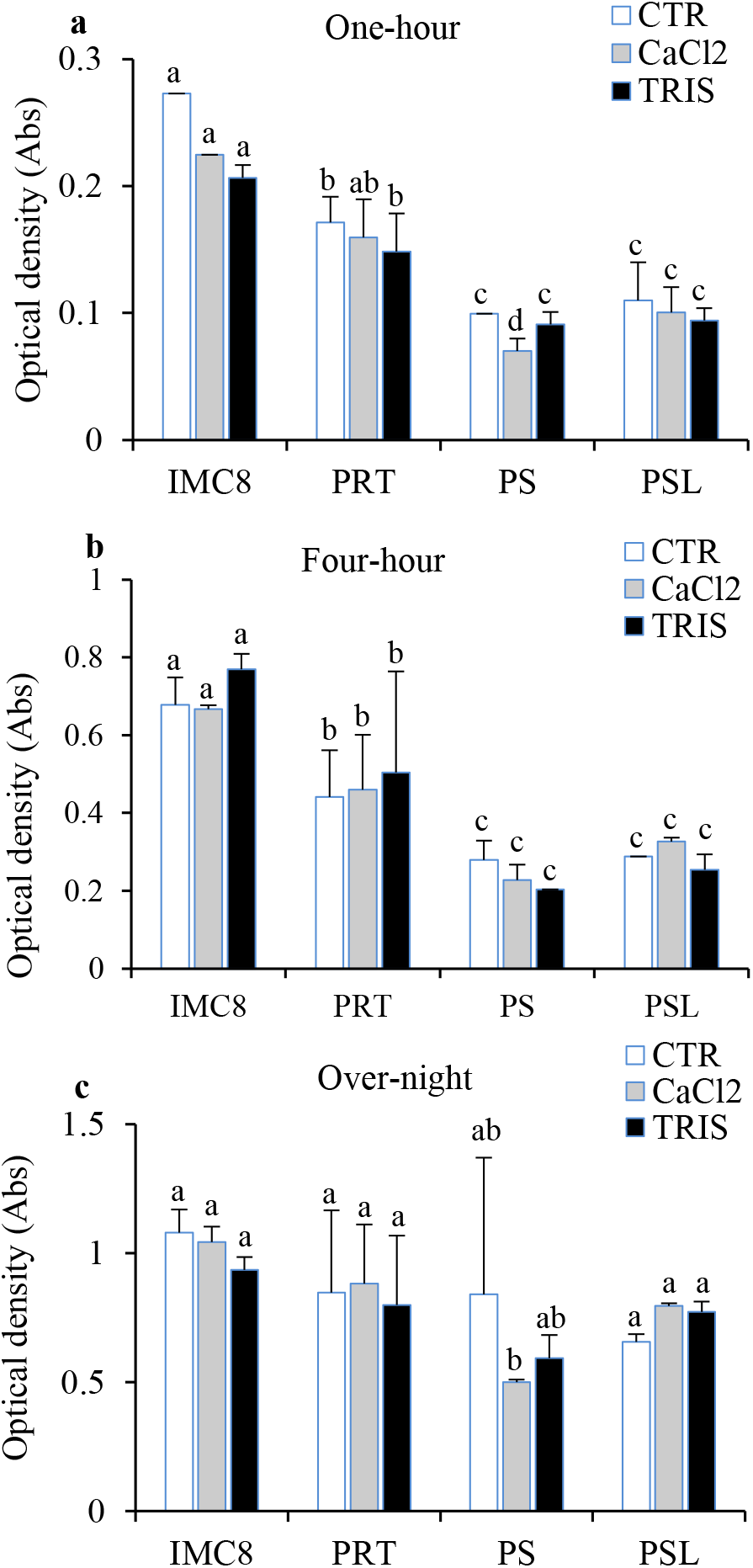
Effects of competency on BCA’s growth. **a** to **c**, Growth characteristics of competent BCA strains IMC8, PRT, PS, and PSL, with control non-treated BCAs (CTR, white bars graph), CaCl_2_-treated BCAs (gray bars graph), and TRIS-treated BCAs (black bars graph). BCA strains grown on LB-Broth for one hour (**a**), four-hour (**b**), and overnight (**c**). The data depicted in the graphs correspond to the mean ± Standard Deviation (SD) of two replications of culture events (n = 3 tubes scored per culture event for each BCA strain).

### Performance of Competent BCAs in Plasmid-Mediated Transformation

Competent BCAs were transformed with plasmids, pBSU101, and pANIC-10A expression vectors containing eGFP, *SPEC*-resistant, *pporRFP, KAN*, and hygromycin B phosphotransferase II (*HYG*) genes, respectively. The procedure was optimized using different plasmids and BCA strains (IMC8, PRT, PS, and PSL) to transform approximatively 0.5 OD_600_ bacterial inoculum. With the pBSU101-mediated CaCl2-treated- and TRIS-treated BCAs transformation, we found that only 1 µl of BCA cells sprayed onto LB plate led to the selection of 4.67±3.79 to 382.33±19.66 and 123.33±18.58 to 393±17.06 spectinomycin-resistance CFU BCAs-eGFP for CaCl2-treated- and TRIS-treated BCAs, respectively (Table 1). Additionally, we found that the transformation efficiencies (TEs) varied significantly from 5.43±2.39% to 58.36±7.78% and from 51.48±4.38% to 83.89±7.41% for CaCl2-treated- and TRIS-treated BCAs, respectively. Similarly, with the pANIC-10A-mediated CaCl2-treated- and TRIS-treated BCAs transformation, we found that up 1.33±0.58 to 2.67±0.58 and 526±99.02 to 763±51.42 kanamycin-resistance CFU BCAs-*pporRFP* for CaCl2-treated- and TRIS-treated BCAs, respectively (Supplementary Table 1). The TEs for CaCl2-treated- and TRIS-treated BCAs varied significantly from 0.99±0.37% to 4.06±0.62% and from 59.46±11.15% to 75.37±6.36%, respectively. Transformation efficiencies varied significantly based on the BCA strain and chemical-induced competency used. The highest TEs were obtained using TRIS-treated BCAs for both plasmids pBSU101 and pANIC-10A. The transformation of TRIS-treated PSL strain was more effective than IMC8 or PS in producing more spectinomycin- and kanamycin-resistant CFUs, with average efficiencies of 83.89 ±741% compared to 51.48 ± 4.38% and 75.37 ± 6.36% compared 59.46 ± 11.17% or 64.6 ± 1.4% in pBSU101 or pANIC-10A plasmids, respectively (Table 1 and Supplementary Table 1). Supporting results obtained with the fluorescence microscope showed that among the BCAs grew on LB plates, all displayed either a bright green or orange/red fluorescence signal compared with the non-transformed control BCAs (Fig. 3a to p1), and this was congruent with our PCR results. The transformability status of BCAs, the stability and expression of reporter genes with both plasmids were confirmed with PCR colony analysis of four individual TRIS-treated BCAs (Tr-BCAs) and non-treated control (NTr-BCAs). The results showed that all the CFU BCAs transformed with either pBSU101 or pANIC-10A contained *eGFP* or *HYG B*, and *pporRFP* amplicons, thus indicating that BCAs were recombinant bacteria (Fig. 3). All control BCAs had no eGFP or pporRFP fluorescence signal or produced PCR amplicons (Fig. 2 and 3).

**Figure 2.**
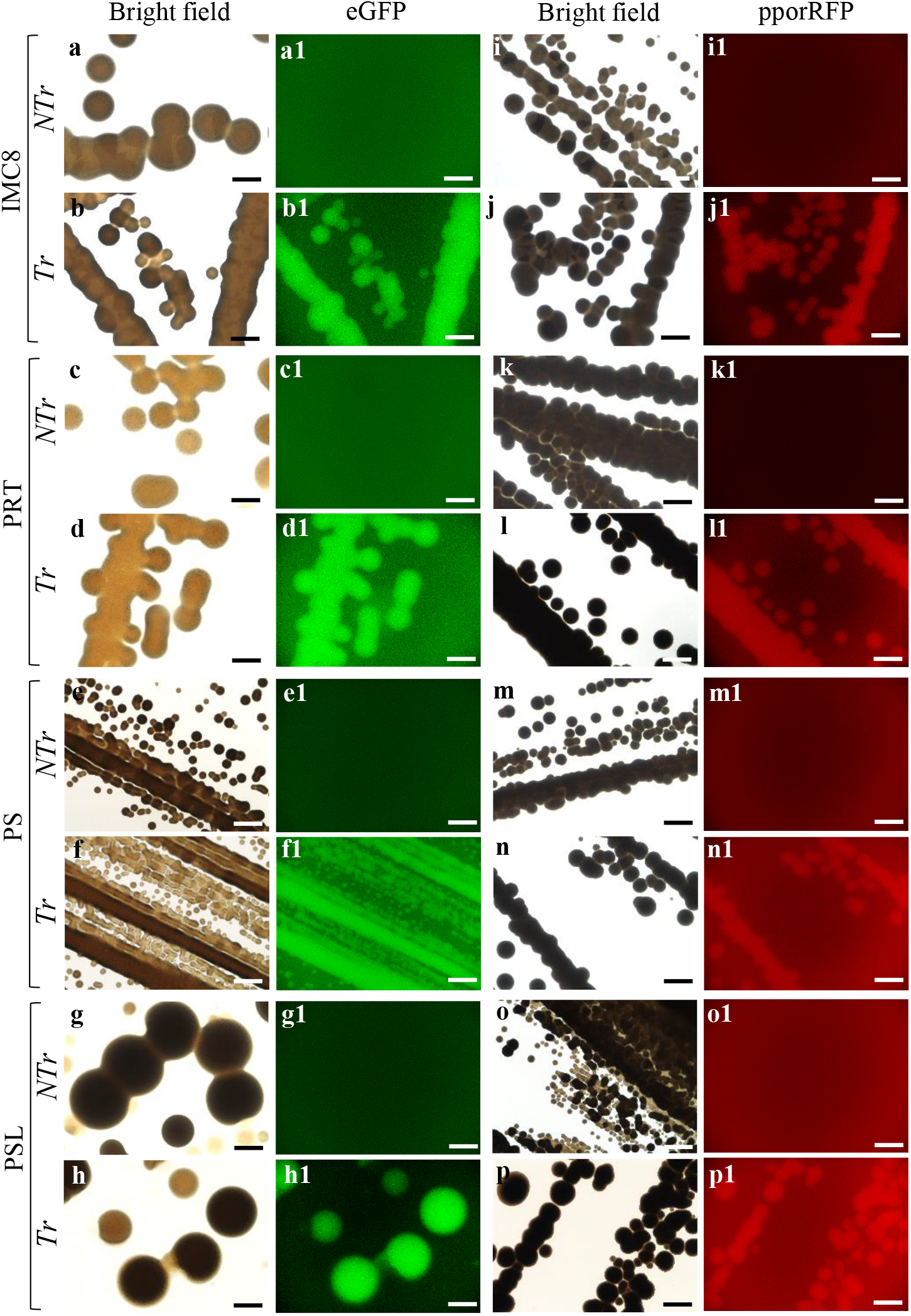
Characteristics of transformed TRIS-treated BCA strains. **a** to **p**, Microscopy micrographs of NTr-BCA and fluorescent Tr-TRIS-treated BCA strains IMC8 (**a** to **b1**), PRT (**c** to **d1**), PS (**e** to **f1**), and PSL (**g** to **h1**) grown overnight expressing green fluorescence of the eGFP. **i** to **p1**, BCA strains IMC8 (**i** to **j1**), PRT (**k** to **l1**), PS (**m** to **n1**), and PSL (**o** to **p1**) expressing orange-red fluorescent of pporRFP protein. **a** to **g**, and **i** to **p** Bright field micrographs. Bars = 400 µm **a** to **p1**.

**Figure 3.**
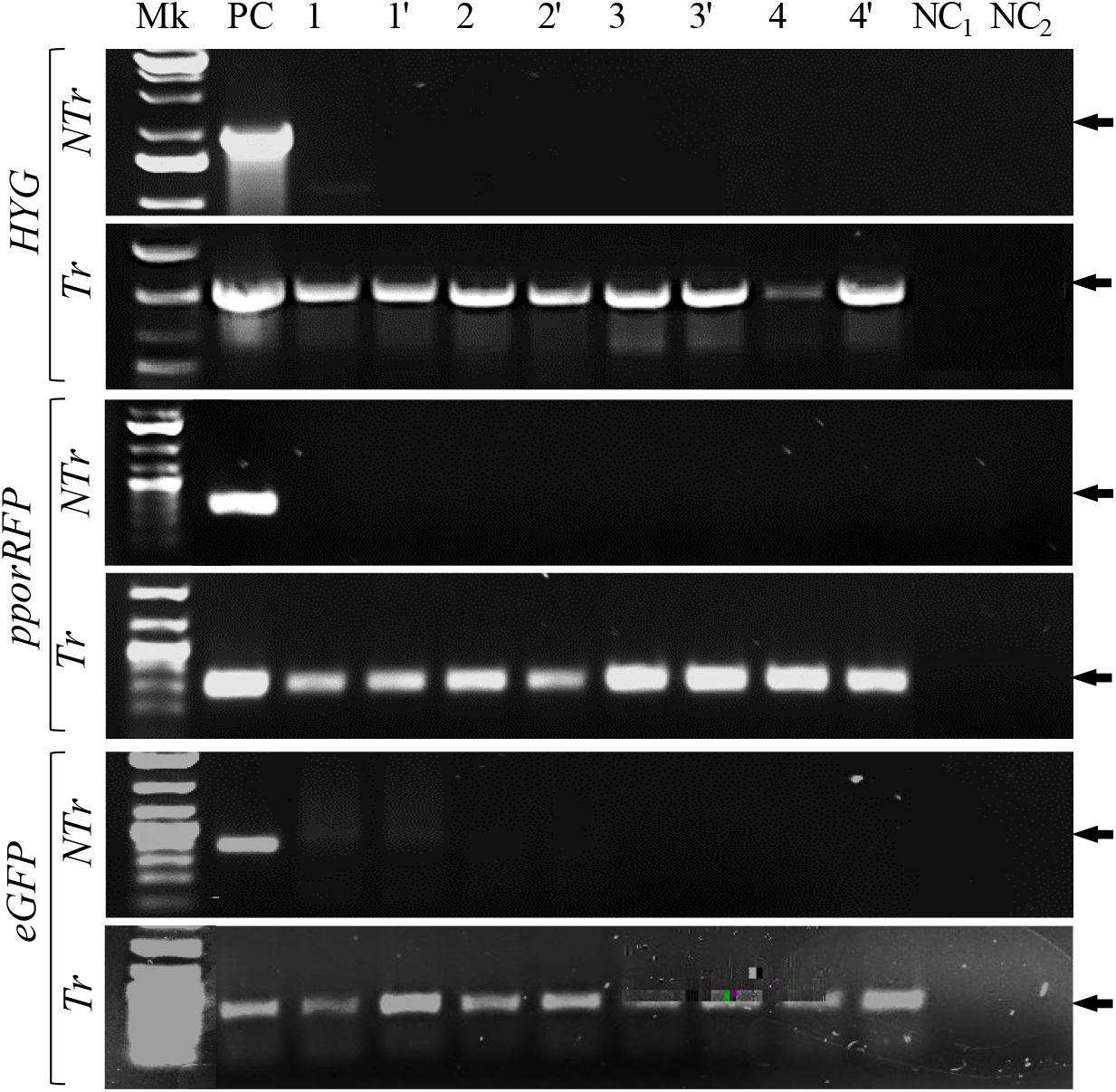
Gene expression of transformed TRIS-treated BCA strains. PCR colony analysis of non-transformed (Ntr-BCA) and transformed (Tr-TRIS-treated BCA) strains to show the presence of *HYG, pporRFP, and eGFP* genes. Mk, Hi-Lo™ DNA marker, PC is positive control of 50 ng of either pANIC-10A (for *HYG* and *pporRFP*) or pBSU101 (for *eGFP*) plasmids DNA. The plasmid DNA (pDNA) extracted from NTr-BCA, and Tr-TRIS-treated BCA strains are two replicates (1-4 and 1’-4’) amplified pDNAs extracted from 2 individual of each BCA strain IMC8 (1 and 1’), PRT (2 and 2’), PS (3 and 3’), and PSL (4 and 4’) respectively. NC_1_ and NC_2_ are negative controls of water (NC_1_) and LB-Broth (NC_2_). Amplicon sizes (indicated by the arrows) are 1 kb, 0.35 kb, and 0.95 kb for *HYG, pporRFP*, and *eGFP* genes, respectively.

Before determining the localization of fluorescent BCAs in the plant, the effect of the process of transformation on either TRIS-treated or CaCl2-treated BCAs IMC8, PRT, PS, and PSL culture was assayed by evaluating growth from frozen bacteria over time. For both types of transformed treated bacteria BCA, the growth on LB antibiotic selection media was well and reproducible. BCAs IMC8, PRT, PS, and PSL strains showed ODs varied between 0.19±0.09 to 0.25±0.16 (for CaCl2-treated BCA) and 0.37±0.25 to 0.55±0.17 OD_600_ (for TRIS-treated) by 1 hr of culture, up to 0.21±0.14 to 0.36±0.23 OD_600_ and 0.23±0.16 to 0.86±0.07 OD by 4 hrs, reaching up to 0.63±0.34 to 0.79±0.28 OD_600_ and 0.69±0.4 to 0.94±0.4 OD_600_ by overnight (Fig 4a, b, c). Both TRIS-treated or CaCl2-treated BCA cell cultures displayed significant densities with a 1.4- to 2.3-fold increase by 4 hrs culture compared to treated BCA controls (CTR-CaCl2 or CTR-TRIS), indicating that the transformation process did not negatively impact the growth of competent BCA strains. However, no significant differences in optical densities were observed for all CaCl2-treated transformed BCAs compared to controls (CTR-CaCl2) for 1 hr culture. Thus, when ODs were compared over time, we determined that BCAs grown for 4 hrs were appropriate inoculums for further plant inoculation assays.

**Figure 4.**
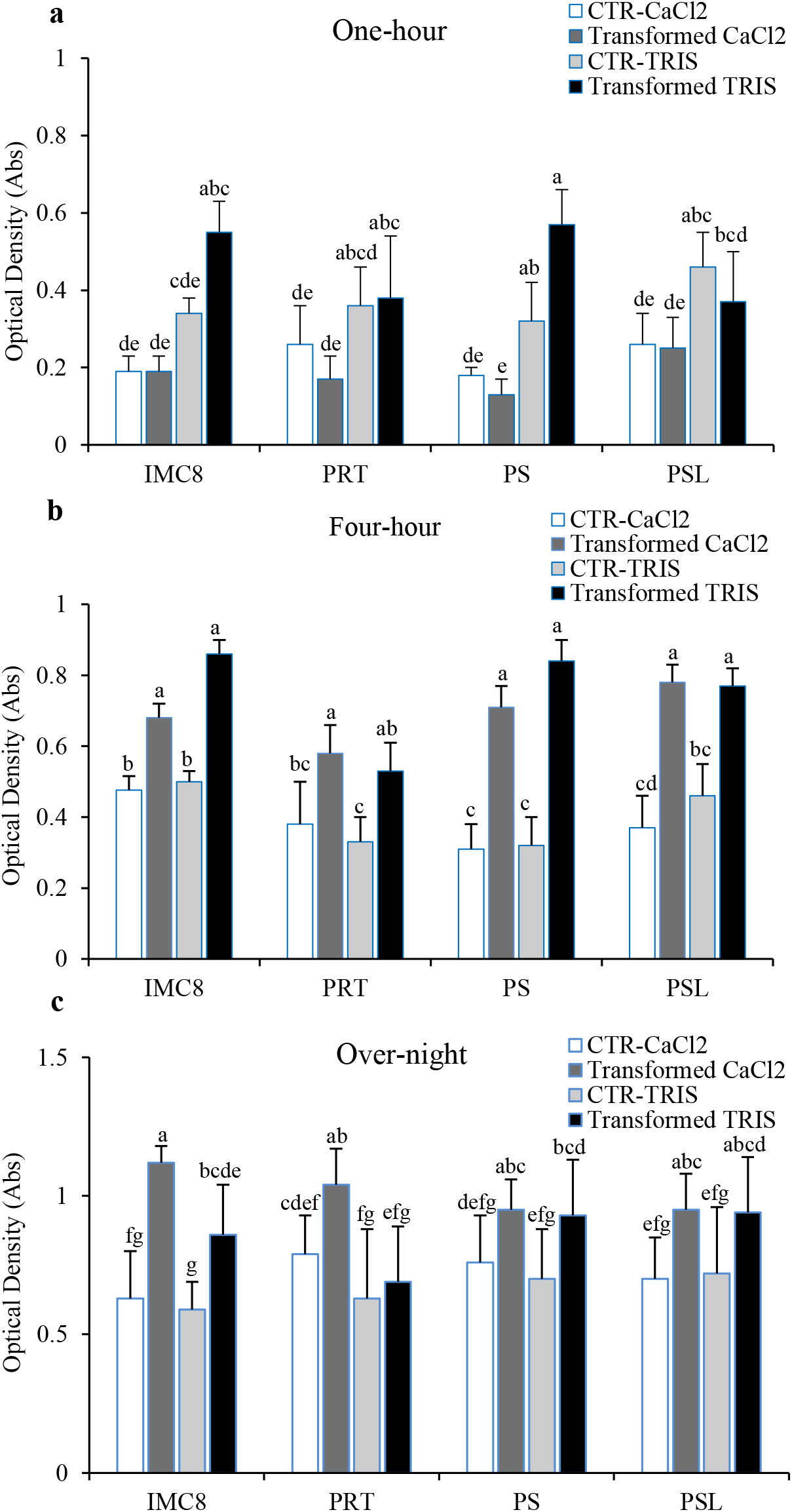
Effects of transformation on BCA’s growth. **a** to **c**, Growth characteristics of plasmid-transformed competent BCA strains IMC8, PRT, PS and PSL with control non-transformed CaCl_2_-treated BCAs (CTR-CaCl_2_, white bars graph), transformed CaCl_2_-treated BCAs (gray bars graph), control non-transformed TRIS-treated BCAs (CTR-TRIS, light gray bars graph), and transformed TRIS-treated BCAs (black bars graph). BCA strains were grown in liquid LB for one hour (**a**), four-hour (**b**), and overnight (**c**), and the growth was evaluated by determining ODs over the time of the culture. The data depicted in the graphs correspond to the mean ± SD of two replications of culture events (n = 3 tubes scored per culture event for each BCA).

Glycerol BCA stocks were used at the start of the assay; therefore, the effects of the cold/freezing on the growth of transformed treated BCAs were evaluated. We found that ODs of either frozen CaCl2-treated or TRIS-treated transformed BCAs were significantly elevated for all treated-transformed BCAs compared to treated-non-transformed BCAs for an overnight culture. With up to 1.99±0.06 to 2.24±0.04 vs. 1.05±0.46 to 1.42±0.31 (for CaCl2-treated transformed and non-transformed BCAs) and up to 1.66±0.02 to 2.23±0.03 vs. 0.69±0.03 to 1.52±0.04 (for TRIS-treated transformed and non-transformed BCAs). The results showed increased rates ranging from 1.40- to 1.60-fold, and 1.3- and 1.7-fold for CaCl2-treated and TRIS-treated transformed BCAs, respectively, compared to treated BCA controls (Supplementary Fig. 1b). These results suggest that the cold or freezing did not negatively impact the growth of recombinant BCA cells.

### Performance of Transformed BCAs in Host-Plant Colonization: Determination of Spatiotemporal Localization of Fluorescent BCAs

Because TRIS-treated BCAs displayed the highest TEs, IMC8-, PRT-, PS- and PSL-tagged to either eGFP- or pporRFP were used for seed inoculums for colonization of Arabidopsis and sorghum plants. The presence of either eGFP or pporRFP fluorescence signals was visualized among five 7- to 14-day-old Arabidopsis or sorghum seedlings. We found that mostly roots or hypocotyls had either bright green eGFP or orange/red pporRFP fluorescent BACs as seen under the FITC or TRITC filter set (Figs. 5a to 5j and 6a to 6h). The spatiotemporal localization of fluorescent BCAs was seen to be associated within all types of root of both plants (Fig. 5a to 5j and Fig. 6a to 6h, and Supplementary videos 1, 2, 3, 4, 5). Fluorescent BCAs were undetectable in non-colonized control seedlings, in roots, leaves for both plants as well as in sorghum stems (Fig. 5a, b and Fig. 6a, b and Supplementary Fig, 2a, b, c). Also, neither eGFP nor pporRFP auto-fluorescence was observed in seedling under the DAPI filter set (Data not shown). It is interesting to note that BCA-eGFPs co-localized with BCA-pporRFPs in the same host-sorghum as we expected them to interact or co-colonize the same plant together in nature (Supplementary videos 5). These anticipated results demonstrate that BCA-mediated colonization is plant tissue-specific and that their spatiotemporal localization is specifically associated with root and hypocotyl tissues.

**Figure 5.**
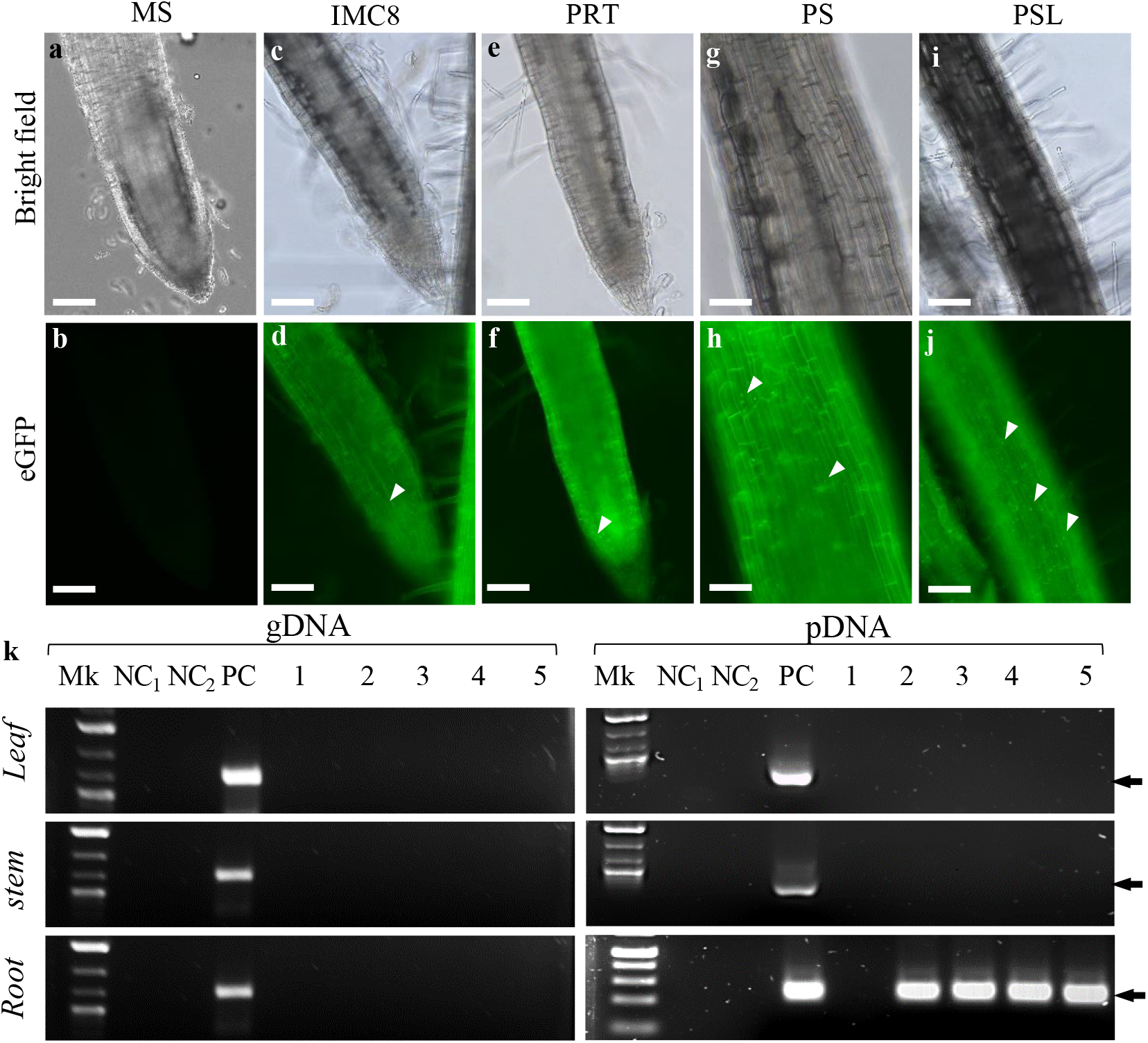
Localization of fluorescent BCAs harboring pBSU101/eGFP in host-sorghum. **a** to **j** Microscopy micrographs of bright field (**a, c, e, g and i**) and fluorescence (**b, d, f, h, and j**) showing fluorescent BCAs expressing eGFP in growing 7-d-old roots of sorghum plants. Non-colonized root, mock ddH_2_O-treated (**a** and **b**); IMC8-pBSU101/eGFP-treated root (**c** and **d**), PRT-pBSU101/eGFP-treated root (**e** to **f**), PS-pBSU101/eGFP-treated root (**g** to **h**) and PSL-pBSU101/eGFP-treated root. The arrowheads indicate BCAs presence. **k**, PCR of gDNA and pDNA DNAs extracted from non-colonized and colonized leaves, stems, and roots showing the presence of the *eGFP* gene. Mk, Hi-Lo™ DNA marker, PC is a positive control of 50 ng of pBS101/eGFP plasmid DNA. The amplicons (1-5) were amplified from gDNA and pDNA extracted from ddH_2_O-(1), IMC8-(2), PRT-(3), PS-(4), and PSL-treated plants (5), respectively. NC_1_ and NC_2_ are negative controls of MS (NC_1_) and LB-Broth (NC_2_). The amplicon *eGFP* size (indicated by the arrows) is 0.850 kb. Bars = 100 µm **a** to **j**.

**Figure 6.**
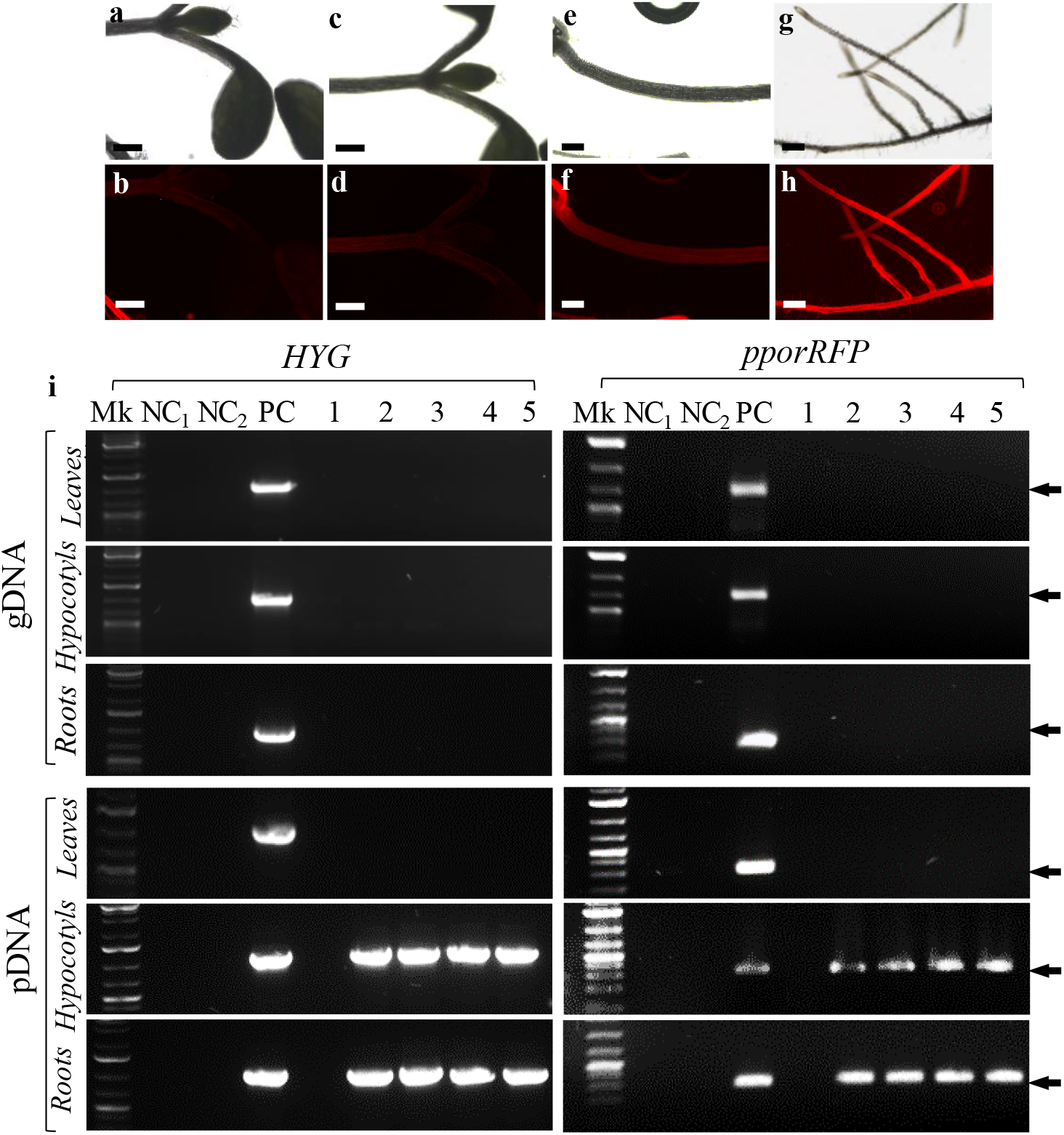
Localization of fluorescent BCAs harboring pANIC-10/pporRFP in host-Arabidopsis. **a** to **j**, Microscopy micrographs of bright field (**a, c, e, g**, and **i**) and fluorescence (**b, d, f, h**, and **j**) showing fluorescent BCAs expressing the pporRFP fluorescent protein (RFP) in 7-d-old Arabidopsis seedlings. Non-colonized shoots, mock ddH_2_O-treated (**a** and **b**); IMC8-pANIC-10A/pporRFP-treated shoot/leaf (**c** and **d**); hypocotyl (**e** and **f**), and roots (**g** and **h**). **i**, PCR of gDNA and pDNA DNAs extracted from non-colonized and colonized leaves, hypocotyls, and roots showing the presence of *HYG* and *pporRFP* genes. Mk, Hi-Lo™ DNA marker, PC is a positive control of 50 ng of pANIC-10A/pporRFP plasmid DNA. The amplicons (1-5) were amplified from gDNA and pDNA extracted from ddH_2_O-(1), IMC8-(2), PRT-(3), PS-(4), and PSL-treated seedlings (5), respectively. NC_1_ and NC_2_ are negative controls of MS (NC_1_) and LB-Broth (NC_2_). The amplicons *HYG* and *pporRFP* sizes (indicated by the arrows) are about 0.900 and 0.350 kb, respectively. Bars = 400 µm **a** to **h**.

In addition to our microscopy observation, the genotype status of the colonized plants, the integration, stability, and expression of inserted reporter genes into colonized plants were investigated using PCR. Analysis of three (3) individual BCA-colonized and non-colonized control plants showed that all roots of both plants and Arabidopsis hypocotyls contained *eGFP*, and both *HYG* and *pporRFP* reporter genes amplicons from only the pDNAs, and not from gDNA indicating that they were not transgenic plants (Fig. 5k and 6i). Amplification of the three reporter gene fragments was not detected in sorghum leaves and stems nor in non-colonized control plants. These PCR results strongly supported our fluorescence localization results obtained with root, hypocotyl, leaf, and stem tissues. All control plants had no evidence of BCA-eGFPs or BCA-pporRFPs fluorescence signal or PCR amplicons.

## DISCUSSIONS AND CONCLUSIONS

### Competency and Transformability of Bacteria *Bacillus*

The generation of electrocompetent bacterial gram-positive strain followed by the heat-shock transformation was reported by Aymanns et al. (2011) and Augustin and Gotz (1990). The authors described a successful construction of EGFP-plasmid as a general tool to transform a large variety of gram-positive species comprising different streptococcus and related genera. However, no further study was conducted on the same gram-positive bacteria using chemically competent cells or the genera *Bacillus*. Our study developed an **e**fficient fluorescence-based localization technique for real-time tracking at single-cell and bulk-level of *Bacillus* biological control agents in host-plants colonization. We first produced competent Bacillus cells using two different methods using Calcium chloride (Chang et al. 2017) and TRIS-HCL (Hofgan and Willmitzer, 1988). Both methods have been used to induce competency, followed by a heat-shock transformation in bacteria gram-negative *Escherichia coli* (Chang et al. 2017) and soil bacterium *Agrobacterium tumefaciens* (Hofgan and Willmitzer, 1988). To perform the plasmid-mediated transformation, the effect of competency on BCA was evaluated to demonstrate that neither Calcium chloride nor TRIS-HCL treatment influenced the growth of competent BCAs compared with non-competent BCAs (Fig. 1 and Supplementary Fig. 1). BCAs transformation performance was tested using two plasmids, pBSU101 (Aymanns et al. 2011) and pANIC-10A (Mann et al. 2012a), that contain reporter genes *eGFP* and *pporRFP*, respectively. This method of transformation also served to reduce the inconsistency of bacteria transformability. We demonstrated that competent BCAs were consistently transformed with plasmid-mediated transformation efficiencies ranging from 51 to 84% (Table 1 and Supplementary Table 2).

Moreover, we were able to produce continuous high-level expression of EGFP consistent with the results reported by Aymanns et al. (2011) for gram-positive bacteria. These findings suggest that the promoter *cfb* to control *egfp* transcription in pBSU101 can also be used as a tool to induce hyperexpression of other genes in gram-positive hosts. Although high transformation efficiencies were obtained with the binary vector pANIC-10A (Supplementary Fig. 1), we observed poor orange/red fluorescent signals from BCA cells or bulks harboring pANIC-10/pporRFP in root tissues of Arabidopsis and sorghum. The low resolution obtained from our epifluorescence microscope is most likely due to the lack of the accurate tdTomato filter set: excitation at 543 nm and fluorescence emission collected from 590 to 610 nm wavelength that is appropriate to visualize the pporRFP protein *in-vivo*. Fluorescence intensity can be evaluated using a spectrofluorometry according to methods described by Millwood et al. (2003); Ondzighi-Assoume et al. (2019) with a Fluorolog^®^-3 system. Overall, this low resolution can be resolved using a scanning confocal microscope with high resolution and equipped with appropriate filter sets.

### Comparative Methods of Bacteria *Bacillus* in Colonization of Host-Plants

Tracking the colonization and localization of the BCA population and their number can be determined with the help of several markers. Previous methods showed that the antibiotic-selection method had been one of the essential markers for studying the infiltration of microorganisms into host plants (Gamalero et al. 2003), but this method was not suitable for tracking microorganism localization. Fluorescent markers are popular for microbial colonization studies (Lu et al. 2004; Cao et al. 2011; Krzyzanowska et al. 2012). In the present context, the most promising technique for identifying microorganisms inside plant hosts is molecular techniques. Fluorescent proteins have been reported as an important tool for identifying prokaryotic organisms (Parker and Bermudez, 1997; Valdivia and Falkow, 1997; Bumann 2001; Poschet et al. 2001). The green fluorescent protein encoded by the GFP gene depends on a marker system to visualize tagged bacteria under a microscope (Tombolini et al. 1997). The use of green fluorescent proteins (GFP) and red fluorescent proteins (RFP) is easier to visualize tagged bacteria under a microscope within the cell as a product of gene expression and is less time-consuming in comparison with the use of dyes or other reporters (Bloemberg, 2007). Although several previous studies (Lu et al. 2004; Bolwerk et al. 2005; Grunewaldt-Stöcker et al. 2007) have demonstrated that fluorescence labeling of endophytes with fluorescence tags contributed to the study on the mechanisms of host-microbe interactions, the localization technique remained unfeasible due to the instability of bacterial populations that was influenced by changing environmental conditions facilitated by plasmids that function as important vehicles (Smalla et al. 2015). Broadly, we demonstrated that plant colonization was improved by using a seed-soaking technique with a reduced inconsistency of plant colonization under controlled *in-vitro* culture conditions and subsequently facilitated BCAs population imaging session. Our protocol is an aseptic technique that uses an optimum seed-soaking approach to ensure biocontrol agent’s uptake to colonize the host plants. Previous studies have proven the effectiveness of the seed-soaking approach under non-sterile conditions for bio-control colonization of several plants, including Cucumber, Cantaloupe, Tomato, Pepper (Abdel-Kader et al. 2012), and in snap beans (Joshua, 2017).

### Limitations of Bacteria *Bacillus* Localization in Colonization of Host-Plants

The technology, as described, is an aseptic procedure but can also be used as a non-aseptic technique. If a more extended imaging period is desired in which infection or contamination may become a factor, consider adopting an aseptic technique. The method uses both an inverted and upright microscope (Keyence BX710 epifluorescence microscope) designed so that the window can be fixed to the microscope stage and thus eliminate tissue movement. However, the upright epifluorescence microscope was the only option available for this study. The significant differences in our fluorescence approach lie in adopting an upright microscope design to secure the specimen to an elevated platform and adaptation of a more flexible temporary imaging chamber. This provides unique strengths in tissue accessibility for fluorescence-guided microscopy of connective fluorescent BCAs. However, one drawback of using an upright epifluorescence microscope is related to the thickness of plant tissue. Thick specimens limit their fixation on the cover slide and subsequently induce damages to tissues if they are force-mounted.

Moreover, the upright microscope used in this study limited the resolution of thin specimens as we saw with Arabidopsis where we were unable to visualize single-cell or bulk-level of bacteria due to the low resolution and missing high magnification objectives of our microscope (Fig. 6a to 6h). The epifluorescence microscope suffers from a low signal-to-noise ratio due to fluorescence above and below the focal plane. Therefore, we suggest strategies to eliminate this issue. The protocol can be implemented on other upright microscopes, including confocal microscopes and a fluorescence stereomicroscope, without the need for pigmenting, but more care will be needed to limit the instability of the host specimen with this technology. The confocal microscope will provide images with enhanced contrast and resolution compared to epifluorescence in thicker samples than the focal plane, allowing optical sectioning (Yao and Carballido-López, 2014). Drifting of thick plant tissues necessitates frequent repositioning of the field during imaging. Breathing movements can affect imaging but can be overcome by adjusting the setup. We provide more detailed steps and considerations for our fluorescent-based BCA localization technique implementation than previous descriptions (Yao and Carballido-López, 2014) so that others with appropriate equipment can use the same approach.

In conclusion, here, we report a novel robust, efficient and reliable fluorescence-based localization technique for real-time temporally spatial-imaging of live bacterial BCAs movement at host plants’ cell, tissue, and bulk levels. This is the first time fluorescent BCAs are produced as a tool to (i) determine their specific locations in colonized plants and subsequently (ii) study microbes and host-plant interactions. This highly fluorescence-localization technique enabled the localization of living bacteria BCAs in two plant hosts in only 1-2 months. Finally, these novel improved tools substantially enhanced plant colonization and interaction potentials in the field, providing a system to study microbe-plant interactions by investigating molecular and physiological mechanisms that underlie these interactions vital for agricultural plants.

## MATERIALS AND METHODS

### Growth and maintenance of bacterial strains

Four selected bacterial BCAs (Supplementary Table 2) in glycerol stock were used to grow fresh cultures on Luria-Bertani (LB) agar for 24 h in a 37°C incubator. Bacterial BCA strains, *Bacillus thuringiencis* (IMC8), *Bacillus subtilis* (PRT), *Bacillus vallismortis* (PS), *Bacillus amyloliquefaciens* (PSL) were streaked onto LB agar plates (LB; 10 g/l Bacto Tryptone, 5 g/l Bacto Yeast, 5 g/l NaCl, and 15 g/l Bacto Agar) to generate colonies of single forming units (CFU) and then harvested and grown for 24h at 200 rpm in 3 ml nutrient broth to generate single-cell cultures. Each single-cell colony culture was used to produce competent BCA cells as described below.

### Generation of competent BCA Cells

Two high-level methods were used: (i) 10 mM Tris-HCl, pH 7.5 (TRIS, Sigma Aldrich) as described by Hofgen and Willmitzer (1988), and (ii) 0.1 M calcium chloride (CaCl_2_, Sigma Aldrich) adapted from Chang et al. (2017). Single-cell colony-forming units of each BCA produced above were grown in 5 ml LB broth and cultured overnight at 37°C on a rotary shaker at 200 rpm. To produce competent BCA cells, 100 µl of each BCA were sub-cultured overnight (with OD_600_ reaching above 0.5 corresponding to approximately 10^8^ CFU/ml). Overnight BCA cultures were pelleted at 3,900 rpm at 4°C for 24 h followed by a treatment of either 10 ml ice-cold 0.1 M CaCl_2_ or 10 mM TRIS-HCl, pH 7.5 solutions for 30 minutes as previously described by Chang et al. (2017) and Hofgen and Willmitzer (1988). Aliquots of 100 µl of generated competent BCA cells were placed in 1.5-ml Eppendorf tubes and stored at -80°C before using them for downstream competency evaluation and transformation.

The evaluation of the grown inoculum placed in 96-well plates was used to determine the original BCA growth by estimating optical densities (ODs) at OD_600_ with a spectrophotometer FLUOstar Omega, BMG Labtech. The original BCA suspension was considered zero time and used as inoculum for BCA competency over time. A dilution of 1:100 from the original BCA was grown in 3 ml LB and cultured for 1 hr, 4 hrs, and overnight. At each time point, titers (optical densities at OD_600_) of competent TRIS-BCA or CaCl_2_-BCA were estimated as described above and compared with the original BCA cultures at zero-time, 1 hr, 4 hrs, and overnight.

### Plasmid-Mediated Heat-Shot Transformation of BCA Strains

The transformation procedure was conducted using a plasmid-mediated DNA delivery method developed by our laboratory modified from previous protocols (Aymanns et al. 2011; Ondzighi-Assoume et al. 2019). Two different plasmids, pBSU101 (Aymanns et al. 2011) and pANIC-10A (Mann et al. 2012a) (listed in Supplementary Table 2) were used to transform both competent TRIS-BCA or CaCl_2_-BCA cell suspension cultures. Both plasmids were constructed in *Escherichia coli*. The plasmid pBSU101 obtained from the Institute of Medical Microbiology and Hygiene, University of Ulm (Ulm, Germany) has an enhanced green fluorescence protein (eGFP) and spectinomycin (*SPEC*) gene that confers resistance to the spectinomycin antibiotic. The binary vector pANIC-10A obtained from the University of Tennessee carries the switchgrass polyubiquitin 1 promoter and intron (*PvUbi1*), which drives the expression of *Porites porites* red fluorescent protein-coding region (*pporRFP*) and hygromycin B phosphotransferase coding region (*HYG*) regulated by switchgrass polyubiquitin 2 promoter and intron (*PvUbi2*) (Mann et al. 2012a; Mann et al. 2012b). The expression of the *pporRFP* serves as a reference of red fluorescence, whereas the *HYG* gene confers resistance to the hygromycin B antibiotic. Before co-cultivation for transformation, 100 µl aliquots of either competent TRIS-BCA or CaCl_2_-BCA suspension were thawed on ice for 30 minutes and then mixed with 0.5-1 µg µl^-1^ of either plasmid pBSU101 or pANIC-10A. Then, either 100 µl mixture TRIS-BCA or CaCl_2_-BCA cells were subjected to the heat-shock method to introduce the plasmid DNA into cells. Briefly, designated mixtures were heat-shocked by placing them in a 42 °C water bath for 45 seconds, transferred on ice for 5 minutes, followed by the addition of 900 µl pre-warmed LB medium. The tubes were kept in a rotary shaker at 37 °C for 1 hr for outgrowth, followed by centrifuging at 10,000 rpm for 3 minutes. Supernatants were discarded, and 200 µl LB was added to the pelleted cells. Separately, two volumes of 50 µl and 150 µl of suspended BCA cells were sprayed onto LB plates containing appropriate antibiotics for each plasmid. BCAs transformed with the plasmid pBSU101 were sprayed onto LB plates contained the antibiotic spectinomycin (*SPEC*^*100*^, 100 µg ml^-1^), and for BCAs transformed with plasmid pANIC-10A, onto LB plates with antibiotic kanamycin (*KAN*^*50*^, 50 µg ml^-1^). After overnight growth, the transformation efficiency was evaluated by scoring the number of CFUs in transformed BCAs (Tr-TRIS-BCA or Tr-CaCl_2_-BCA) on LB plates without antibiotics compared with the non-transformed BCA on LB plates containing appropriate antibiotics. The transformation efficiency was calculated as follows:

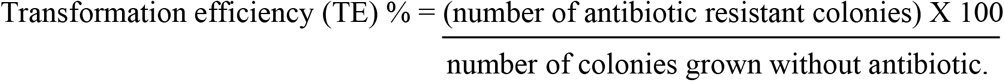

### Host-plant inoculation and growth conditions

*Arabidopsis thaliana* genotype Columbia (Col-0) accession and sweet sorghum variety Topper 76-6 were used as plant hosts for this study. *Arabidopsis* Col-0 seeds were surface-sterilized using 10% ethanol for 10 minutes and then with 10% bleach for 20 minutes. Seeds were rinsed with sterilized distilled water, blot-dried with heat sterilized tissue paper, and then soak-treated in BCA suspensions at 4 °C for 24 hrs. BCA treatments were done with strains IMC8, PRT, PS, PSL, and sterile water (as a negative control). Each BCA inoculum concentration was adjusted to 0.5 at OD_600_. Sorghum Topper 76-6 seeds were surface sterilized using 75% Clorox bleach for 1 hr in a rotary shaker at a speed of 400 rpm. Seeds were thoroughly rinsed in sterile ddH_2_O before inoculation with BCAs, as explained above. Inoculation time of seeds with BCAs was one hour in a shaker at 400 rpm at room temperature. Seeds were then dried with a sterile filter paper and planted on solid half-strength (1/2 X) MS (Murashige and Skoog, 1962) in sterile Petri dishes for Arabidopsis or sterile magenta boxes for sweet sorghum seeds. Either magenta boxes or plates were sealed with Parafilm and placed vertically in a Caron MTR30 growth chamber under a 16-h-light/8-h-dark photoperiod with a light intensity of 100µmol m22s 21 at 22°C, with a relative humidity level between 70-80%.

### DNA Isolation, PCR and Gel Electrophoresis Assays

PCR analysis was used as previously described by Ondzighi-Assoume et al. (2016, 2019) to confirm transformation events of the BCAs and plant colonization. Plasmid DNA (pDNA) was isolated from a single BCA colony, leaf, hypocotyl, and root tissues using a Qiaprep Spin Miniprep kit (Qiagen). At 14 d (16 days post-inoculation, DPI), pDNA and genomic DNA (gDNA) were isolated from plant tissues as previously described by Edwards et al. (1990). Polymerase chain reaction (PCR) assay was performed to confirm the presence of expressed marker genes in transformed BCAs strain, and BCA colonized plants. To determine the presence of transformed BCA strains from the BCA population living out or inside the plant, PCR reactions were performed using EconoTaq Plus Green 2X Master Mix with the Eppendorf Master Cycler ProS. Marker genes *HYG, eGFP*, and *pporRFP* were amplified using established primer sets (Supplemental data: Table S2). The primer concentration used was 0.1 µM for each forward and reverse primer. The annealing temperature (Tm) was adjusted (+3 or -3 °C) according to the Tm of each primer. PCR was set at a standard 30-cycle reaction with denaturation, annealing, and extension temperatures of 94°C, 53°C, and 72°C, respectively. PCR products were verified with gel electrophoresis using 1% agarose LE (BioExell).

### Fluorescence Microscopy

The analysis of transformed BCA colonies or BCA-inoculated Arabidopsis or sorghum plants was performed as previously described (Ondzighi-Assoume et al. 2019). Either transformed bacterial BCAs or inoculated seedlings were observed with 4x objective in Keyence BX710 epifluorescence microscope (www.Keyence.com)using either the FITC filter set with an excitation wavelength (Ex) at 490 nm and emission detection (Em) at 525 nm (<u>Fluorescence -Flow Cytometry Guide | Bio-Rad (bio-rad-antibodies.com</u>) for GFP or TRITC filter set with an excitation wavelength (Ex) at 544 nm and an emission wavelength (Em) at 570 nm (<u>Tetramethylrhodamine (TRITC) | Thermo Fisher Scientific - US</u>) to visualize pporRFP. LB plates containing transformed BCA colonies were placed under the microscope. Arabidopsis and sorghum seedlings at either 7-d (9 DPI) or 14-d (16 DPI) were mounted on a microscope slide, covered with a cover slide, and visualized under the fluorescent microscope Keyence to check for either EGFP or pporRFP fluorescence signal. The controls were also visualized under the fluorescent microscope to confirm the absence of fluorescence in non-transformed BCA and non-infected plants. The acquisition of videos of BCAs movement was performed using an iPhone from the computer screen.

### Statistical Analysis

Experimental data obtained in this study were generated with PROC GLM, SAS v. 9.4. Analysis of BCAs growth characterization and transformation efficiency data, significance of differences between different treatment groups was assessed using analysis of variance (ANOVA). The specific difference in mean was compared using the least significant difference test (LSD) at P ≤ 0.05, and treatments are presented as mean ± standard deviation (SD). All experiments in this study were performed in triplicate. According to SAS analysis, different letters denote a statistically significant difference among means at a p-value < 0.05 for all the figures and tables presented in this work.

## ABBREVIATIONS

BCA: Biological Control Agents
CaCl_2_: Calcium Chloride
CFU: Colony Forming Unit
ddH_2_O: Double Distillated water
EGFP: Enhanced Green Fluorescent Protein
IMC8: *Bacillus thuringiensis*
MS: Murashige and Skoog medium
gDNA: Genomic DNA
LB: Luria-Bertani
OD: Optical Density
pDNA: Plasmid DNA
PRT: *Bacillus subtilis*
PS: *Bacillus vallismortis*
PSL: *Bacillus amyloliquefaciens*
RFP: Red Fluorescent Protein
TAE: Tris Acetate-EDTA [Ethylenediamine-Tetraacetic Acid] buffer
TE: Transformation Efficiency
TRIS: Tris-based solution

## SUPPLEMENTARY INFORMATION

**Supplementary Figure 1**. Effects of the freezing on competent and transformed BCA’s growth. **a**, Growth characteristics of competent BCAs with control non-treated (CTR, white columns graph), CaCl_2_-treated (gray columns graph), and TRIS-treated (black columns graph) BCAs. **b**, Growth characteristics of transformed competent BCAs with control non-transformed competent CaCl_2_-treated (CTR-CaCl_2_ white columns graph), transformed competent CaCl_2_-treated (gray bars graph), control non-transformed competent TRIS-treated (CTR-TRIS, light gray bars graph), and transformed competent TRIS-treated (black bars graph) BCAs. The data depicted in the graphs correspond to the mean ± SD of two replications of culture events (n = 3 plates scored per culture event for each BCA strain).

**Supplementary Table 1**. Transformation efficiencies of the pANIC-10A-mediated transformation of BCA cells.

The numbers represent the efficiencies of the plasmid pANIC-10A transformation of BCA strain IMC8, PRT, PS, and PSL cultures. The CaCl_2_- or TRIS-induced competent BCA cells were transformed simultaneously at the final concentration of inoculum OD_600_ = 0.5. The transformation efficiency was evaluated by scoring a growing kanamycin-resistant (*KAN*^*R*^) colony-forming unit (CFU). The data depicted in the table corresponds to the mean ± SD of two replications of transformation events (n = 3 plates scored per transformation event for each BCA strain).

**Supplementary Figure 2**. Co-localization of PSL-pBSU101/eGFP and IMC8-pANIC-10A/pporRFP populations within sorghum plants.

**a** and **b**, 7-d-old co-colonized sorghum Dale (**a**) and Topper 76-6 (**b**) growing on MS. **c** to **e**, Microscopy micrographs of fluorescent PSL-pBSU101/eGFP and IMC8-pANIC-10A/pporRFP populations in stems. **f**, Green fluorescent PSL-pBSU101/eGFP population in the root. **g**, Red fluorescent IMC8-pANIC-10A/pporRFP population in the root. **h**, overlapped micrograph showing green PSL-pBSU101/eGFP and red IMC8-pANIC-10A/pporRFP populations in the same root. **g**, 7-d-old co-colonized sorghum plants. Bars = 200 µm **c** to **h**.

**Supplementary Table 2**. Selected BCA strains characteristics.

**Supplementary Table 3**. Plasmids and sequences of *HYG, eGFP*, and *pporRFP* gene primers used for PCR.

**Supplementary Video 1**. Movement of IMC8-pBSU101/eGFP population within 7-d-old colonized sorghum root. The video was recorded with a smart iPhone from the visualization on the computer monitor projected by the microscope Keyence.

**Supplementary Video 2**. Movement of PRT-pBSU101/eGFP population within 7-d-old colonized sorghum root. The video was recorded with a smartphone (iPhone) from the computer monitor projected by the microscope Keyence.

**Supplementary Video 3**. Movement of PS-pBSU101/eGFP population within 7-d-old colonized sorghum root. The video was recorded with a smartphone (iPhone) from the computer monitor projected by the microscope Keyence.

**Supplementary Video 4**. Movement of PSL-pBSU101/eGFP population within 7-d-old colonized sorghum root. The video was recorded with a smart iPhone from the visualization on the computer monitor projected by the microscope Keyence.

**Supplementary Video 5**. Co-localization of PSL-pBSU101/eGFP and IMC8-pANIC-10A/pporRFP populations within 7-d-old co-colonized sorghum Topper 76-6 root.

Movement of PSL-pBSU101/eGFP (in green) and IMC8-pANIC-10A/pporRFP (in orange-red) populations in the same root. The video was recorded with a smart iPhone from the computer monitor projecting the movement of BCAs from the microscope Keyence.

## AUTHOR CONTRIBUTIONS STATEMENTS

COA conceptualized the research project, developed, designed, and carried out inoculation procedure, microscopy experiments, data analysis, prepared the figures, and wrote the manuscript. BB performed BCAs competency, transformation experiments, PCR assays, gels electrophoresis, and statistical analyses. AT carried out competent BCAs growth characteristics. ES helped with the maintenance of BCA strain cultures. WKO contributed to perform experiments related to the establishment of competent BCA strains. MM worked on and improved the manuscript. All authors read and approved the final manuscript.

## ACKNOWLEDGEMENTS

The authors would like to thank and acknowledge Ms. Jamille Y. Robinson and Gabriel Omar Carrasquillo for their help in providing inputs on the manuscript. We are thankful to Dr. Barbara Spellerberg from Institute of Medical Microbiology and Hygiene, University of Ulm, Germany, for graciously sending us the plasmid pBSU101. We also thank Dr. Neal Jr. Stewart from The University of Tennessee, Knoxville, for generously giving us the plasmid pANIC-10A.

## COMPETING INTERESTS

The authors declare no competitive interests.

## Notes

**Finding Information:** This project is supported by USDA-NIFA Evans Allen fund no. TENX-1710-SE and USDA Grant Award no. 2017-38821-26418 (TENX-2016-06533).

